# Vividness of Visual Imagery Supported by Intrinsic Structural-Functional Brain Network Dynamics

**DOI:** 10.1101/2024.03.02.582470

**Authors:** Timo L. Kvamme, Massimo Lumaca, Blanka Zana, Dunja Paunovic, Juha Silvanto, Kristian Sandberg

## Abstract

Vividness of visual imagery is subject to individual variability, a phenomenon with largely unexplored neurobiological underpinnings. By analyzing data from 273 participants we explored the link between the structural-functional organization of brain connectomes and the reported intensity of visual imagery (measured with VVIQ-2). Employing graph theory analyses we investigated both the structural (DTI) and functional (rs-fMRI) connectomes within a network of regions often implicated in visual imagery. Our results indicate a relationship between increased local efficiency and clustering coefficients in the structural connectome in individuals who experience more vivid visual imagery. Increased local efficiency and clustering coefficients were mirrored in the functional connectome with increases in left inferior temporal regions, a region frequently identified as a critical hub in the visual imagery literature. Furthermore, individuals with more vivid imagery were found to have lower levels of global efficiency in their functional connectome. We propose that the clarity and intensity of visual imagery are optimized by a network organization characterized by heightened localized information transfer and interconnectedness. Conversely, an excessively globally integrated network might dilute the specific neural activity crucial for generating vivid visual images, leading to less locally concentrated resource allocation in key regions involved in visual imagery vividness.

## 1. Introduction

Visual imagery involves visualizing images in our “mind’s eye” in the absence of external stimuli. It plays a role in various cognitive processes such as memory, emotions, and creativity (Dijkstra et al., 2019; Floridou et al., 2022; Kosslyn et al., 2001; Pearson, 2019; Pearson et al., 2015). Additionally, imagery is widely used in clinical interventions to improve mental health outcomes (Skottnik and Linden, 2019) and has even allowed patients in vegetative states to communicate with the outside world (Monti et al., 2010; Owen et al., 2006). A neuroscientific understanding of imagery is thus of both basic science and clinical relevance, and it has received a surge of interest in recent years (Dijkstra et al., 2022, 2019; Dijkstra and Fleming, 2023).

Much of the recent research on imagery has focused on various aspects of its functional characteristics, comparing, for example, the spatial and temporal overlap between imagery and perceptual processing (Dijkstra et al., 2018; Nadine Dijkstra et al., 2017). A view has emerged that imagery relies on similar cortical areas as perception, but without initial feedforward activity, with greater similarity in high-level areas (particularly when details are not required) and with greater top-down functional connectivity (Dijkstra, 2024; N. Dijkstra et al., 2017). Examination of the structural aspects of the brain enabling imagery has received less attention.

From a network neuroscience perspective, the brain may be viewed as a connectome: a network of nodes as individual neural populations that are linked by edges as axonal pathways for structural connectomes or signaling pathways for functional connectomes (Lumaca et al., n.d.). In the current study, we relate individual differences in imagery vividness to properties of structural and functional connectomes within the context of graph theory.

Building on the results of connectivity studies, evidence from lesion studies, meta-analytic fMRI-task-based activation studies, and systematic reviews, we identified regions of interest constituting a hypothesized “Visual imagery Network”. This network encompasses bilateral regions of the inferior frontal gyrus (specifically the orbital, triangular, and opercular areas) (N. Dijkstra et al., 2017; Winlove et al., 2018), the parahippocampal gyrus (Fulford et al., 2018; Spagna et al., 2021; Tullo et al., 2022), the inferior temporal gyrus and sulcus, lateral occipital temporal sulcus and gyrus (fusiform) (Bartolomeo, 2002; N. Dijkstra et al., 2017; Fulford et al., 2018; Haxby et al., 2001; Ishai et al., 2000; Spagna et al., 2021; Winlove et al., 2018), intraparietal sulcus (N. Dijkstra et al., 2017), precuneus (Cavanna and Trimble, 2006; Knauff et al., 2003), and the occipital pole (N. Dijkstra et al., 2017) (as defined by the Destrieux Parcellation atlas) (Destrieux et al., 2010) (see supplementary material for details).

Examination of this network allows us, for the first time, to probe the topological organization of the brain network related to individual differences in visual imagery vividness. Local efficiency provides insights into processing abilities within interconnected groups of brain regions reflecting functional segregation in the brain (Iturria-Medina et al., 2008). The clustering coefficient reveals how connected certain neural regions are, on a level indicating their interconnectivity strength (Long et al., 2023). Global efficiency provides insight into how the network can process information in parallel, which shows how well the brain integrates its functions (de Pasquale et al., 2016; van den Heuvel et al., 2009). Lastly, betweenness centrality helps us understand the nodes that play a role in controlling and directing the flow of information across the network. We assess both network-level measures that give a view of the overall “macroscopic view” of the structure and efficiency of the entire network, while node-level measures focus on the “microstructure view” providing characteristics of individual nodes within the network. By considering all these measurements we can gain an understanding of how the brain’s connectivity structure relates to different levels of vividness experienced during visual imagery. A Study overview is presented in Fig. 1.

**Figure 1.**
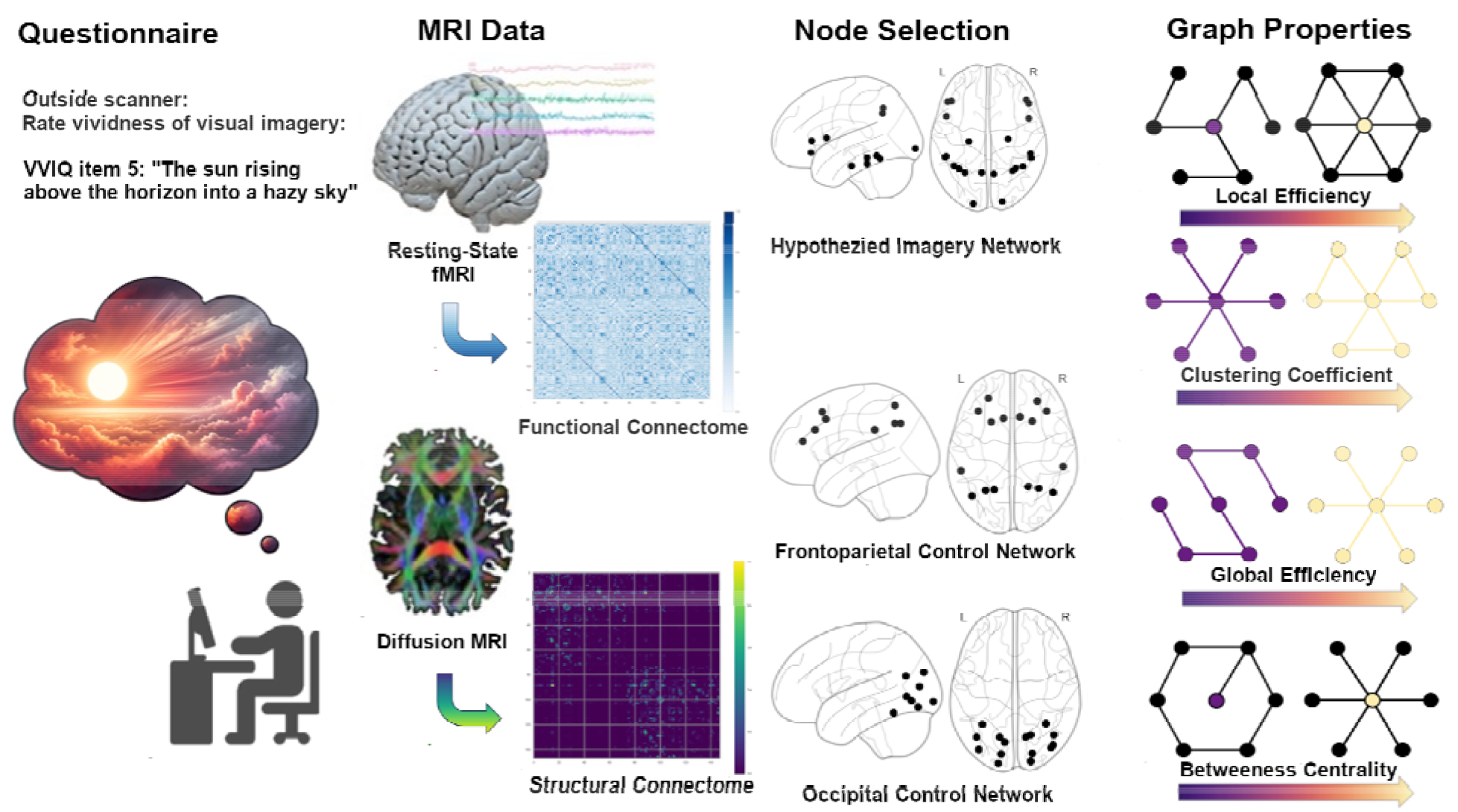
Study overview: Participants rated their visual imagery vividness (VVIQ-2) outside the scanner. We collected resting-state fMRI and diffusion MRI measures to construct structural and functional connectivity matrices. We selected nodes in a hypothesized visual imagery network based on prior task-fMRI studies and tested against two control networks in the frontoparietal and occipital regions. Graph properties of the network were calculated and evaluated using multiple linear regression.

## 2. Results

### 2.1 VVIQ distribution

The participants (n=273) had on average a VVIQ of 0.517 (SD=0.131) on our linear transformed measure of summed VVIQ scores (see Fig 2). The distribution was non-normally distributed (Shapiro-Wilk test, W = 0.988, p=0.02) with a unimodal peak. We found that VVIQ did not differ between males and females (p=0.97) but was borderline significantly correlated with age R = 0.113 (p = 0.06) (see supplementary code for details). All further analyses are corrected for age and gender.

**Figure 2.**
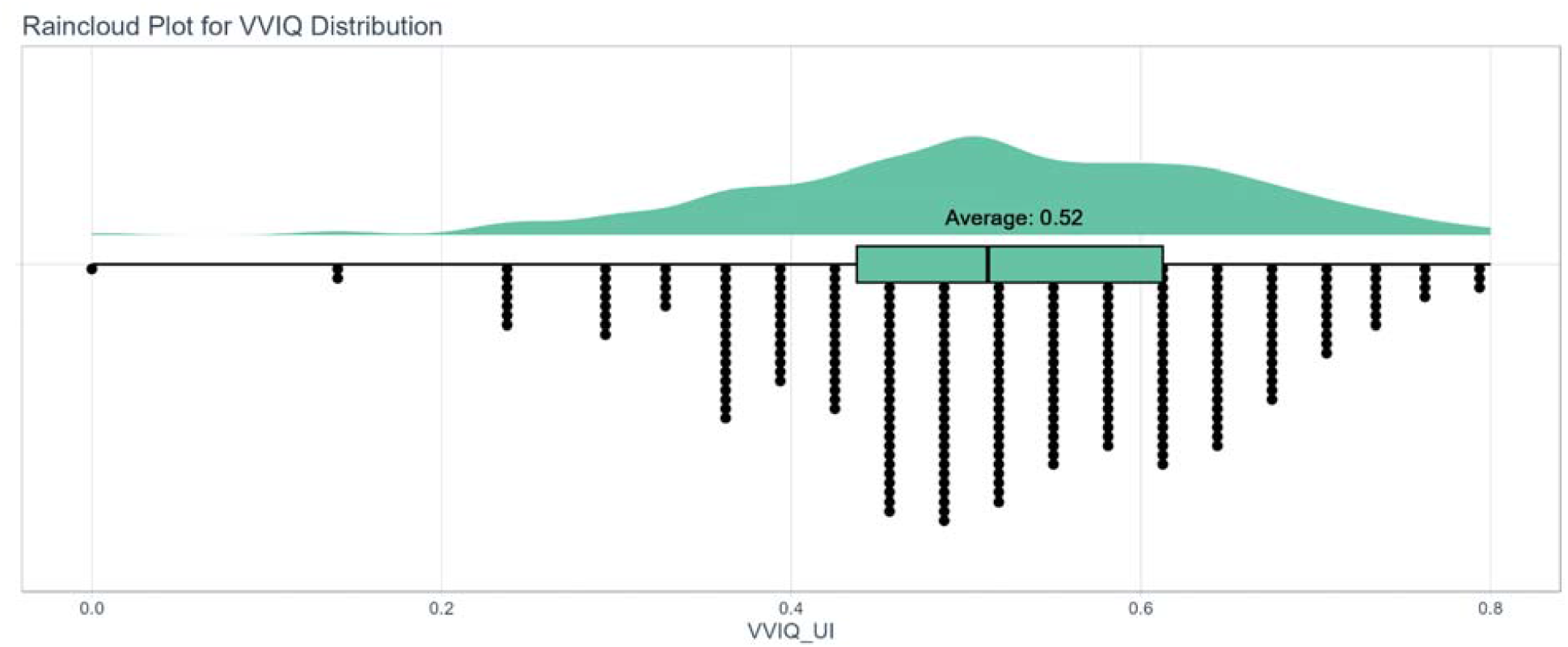
Plot of VVIQ Distribution.

### 2.2 Structural (DTI) Connectome

Two network-level effects were found for the structural connectome (DTI) in the hypothesized visual imagery network for local efficiency and clustering coefficient.

Local efficiency correlated with VVIQ, R=0.170, p=0.007 (see Fig. 3A). For the same network-level structural connectome, we found an R=0.153, p=0.013 for the clustering coefficient (see Fig. 3C). These findings suggest that individuals with higher local efficiency and clustering coefficient in their structural connectome of the hypothesized visual imagery network experience more vivid visual imagery as measured by VVIQ.

**Figure 3.**
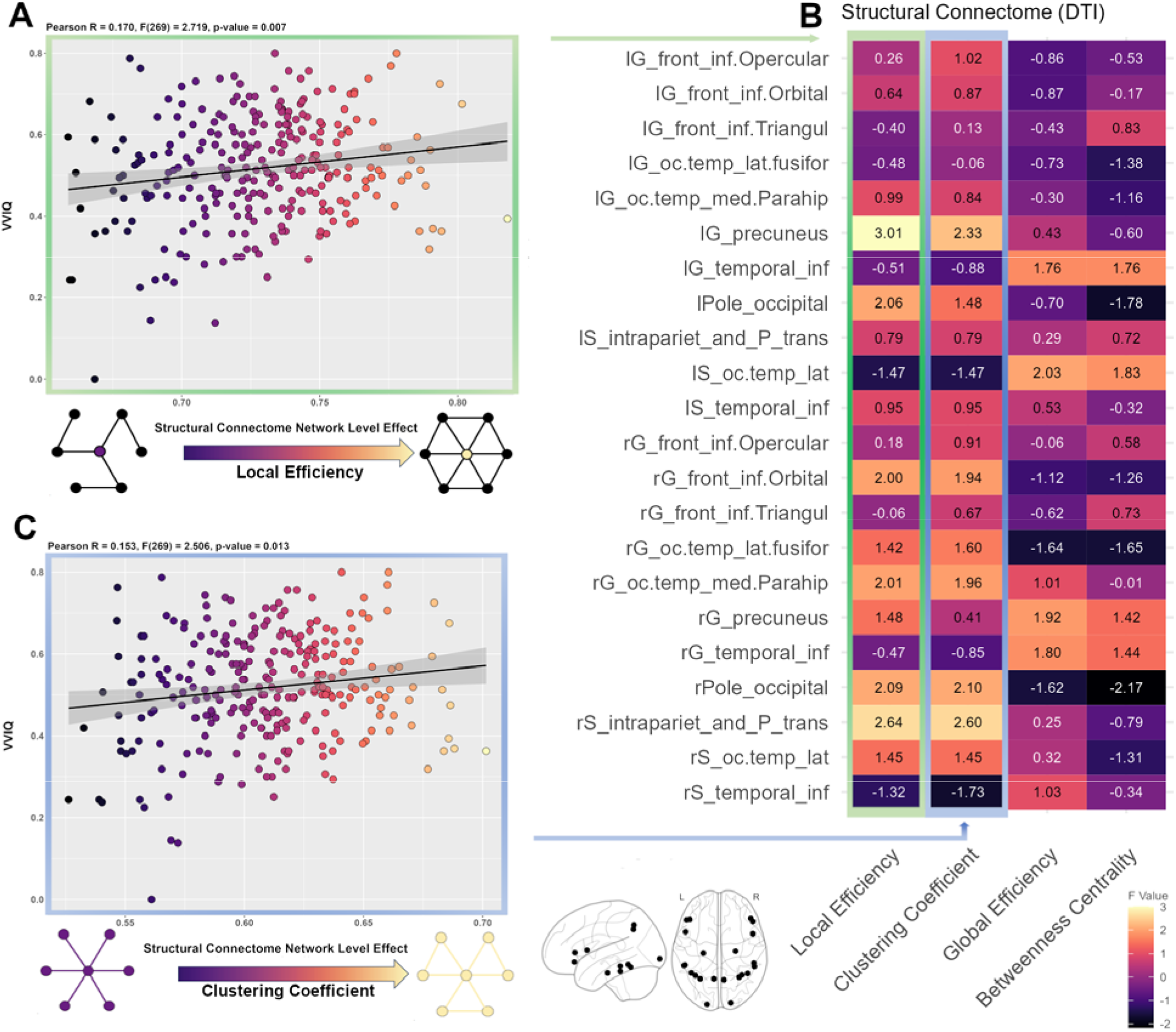
Results for structural connectome (DIT).

No node-level (region) effects were found for the structural connectomes (see Fig. 3B for a table of F values for nodes). However, we observed greater numerical values for local efficiency and clustering coefficient for the left precuneus, the right intraparietal sulcus, and transverse parietal sulci indicating that they are contributing to the significant effect at the network level. Performing the same analysis with the two different control networks of the frontoparietal and occipital structural connectome networks did not result in any correlation that reached significance for either network or node-level effects (see supplementary information).

### Functional (rs-fMRI) Connectome

A significant network-level effect was observed in the functional connectome (rs-fMRI) in the hypothesized network which related to the global efficiency of the network. A negative correlation between network-level global efficiency and VVIQ R = -0.113, p=0.046 was observed (see Fig. 4A).

**Figure 4.**
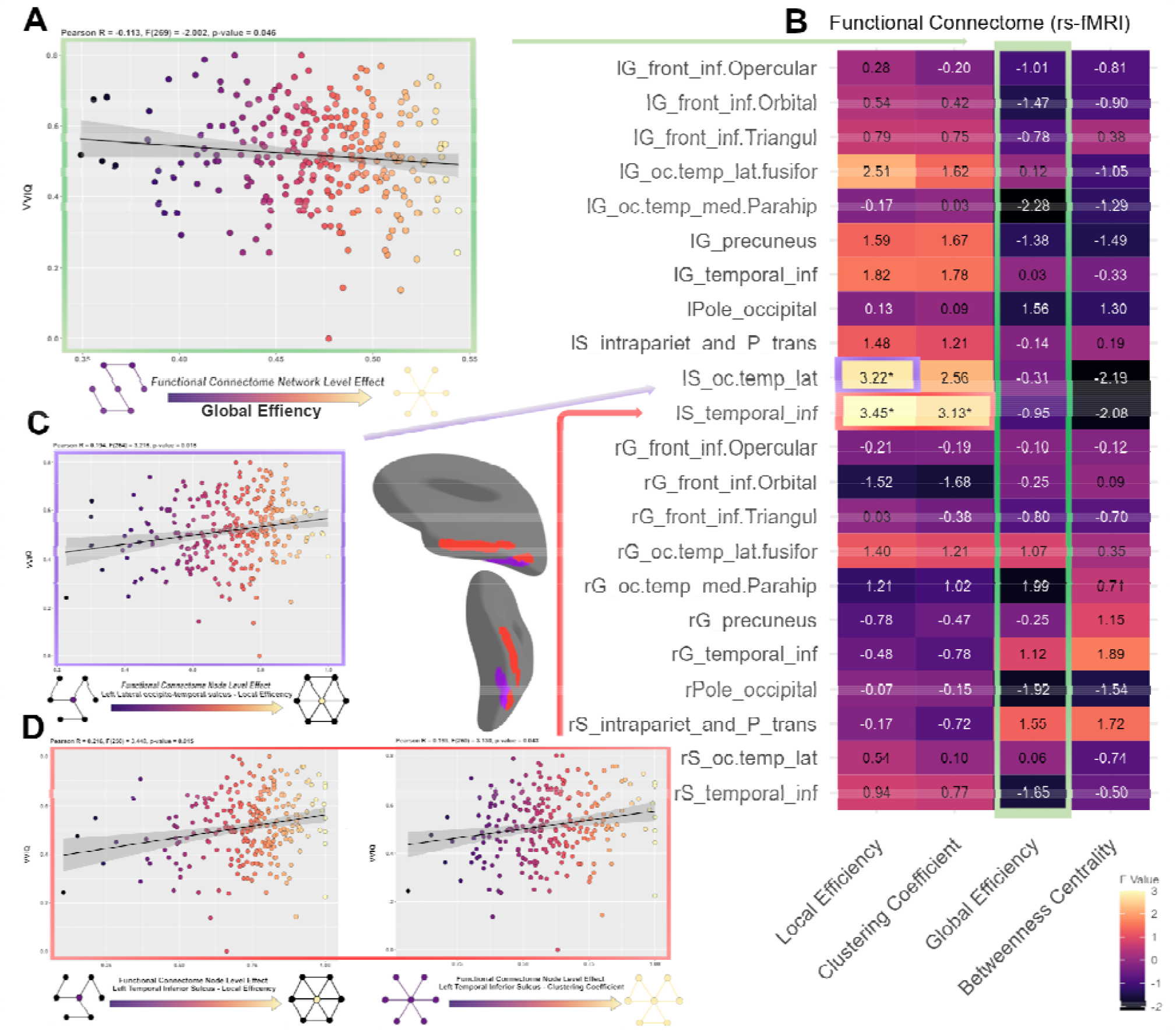
Results for functional connectome (rs-fMRI).

Significant node-level effects were found in the functional connectome with the local efficiency in the left lateral occipitotemporal sulcus correlating with VVIQ with an R = 0.194, p=0.016 (see Fig. 4C). Additionally, the left temporal inferior sulcus had a local efficiency correlation of R = 0.216, p=0.015 with VVIQ, while the clustering coefficient in the same left temporal inferior sulcus region demonstrated a correlation of R = 0.193, p=0.043 with VVIQ (see Fig. 4D). See Fig. 4B for a list of F-values for all regions.

These findings indicate that individuals with lower global efficiency in the functional connectome of the hypothesized visual imagery network, experience more vivid visual imagery. In addition, the findings demonstrate that enhanced local efficiency in the left lateral occipitotemporal sulcus along with enhanced local efficiency and clustering coefficients in the left temporal inferior sulcus are linked to increased vividness of visual imagery. The results underscore the important roles played by these brain regions in the intensity of imagery as measured by the VVIQ.

Performing the same analysis with the two different control networks of the frontoparietal and occipital structural connectome networks did not result in statistically significant effects at the network or node level (see supplementary information).

## 3. Discussion

Our study has illuminated the intricate relationship between the brain’s structural and functional connectomes and self-reported vividness of visual imagery using the VVIQ-2 in a sample of (n=273) participants. By employing graph theory analyses of structural-functional connectomes, we have uncovered significant findings at both the network and node levels of a hypothesized network supporting visual imagery vividness.

At the network level in the structural connectome, we observed that local efficiency and clustering coefficients are positively associated with the ability to experience more vivid visual imagery. This suggests that a more interconnected and efficient local brain network, particularly in areas hypothesized to be part of the visual imagery network, is related to the vividness of mental images. Local efficiency, a measure of the efficiency of information processing within localized brain regions, reflects the network’s fault tolerance. Calculated as the inverse of the average shortest path length among a node’s neighbors, it shows how effectively information is processed locally, even when some nodes or connections fail (Sporns, 2011; Wang et al., 2018; Wig, 2017). Higher local efficiency signifies a network’s ability to maintain functionality despite partial disruptions, demonstrating robust fault tolerance.

Similarly, the observed higher clustering coefficients in the structural connectome indicate that brain regions involved in the hypothesized visual imagery network tend to form more tightly knit groups or clusters in individuals with higher VVIQ scores.

One interpretation is that vivid visual imagery is enabled by a structural network where nodes are highly interconnected (high clustering); furthermore, the network tends to be more efficient locally because there are multiple pathways for information flow within clusters. The combination of efficient local processing and dense interconnections within these clusters appear to facilitate the generation of more vivid visual imagery.

Although we did not observe any significant node-level effects for structural connectome, we saw that the highest F scores were for the left precuneus and right intraparietal and transverse parietal sulcus for the local efficiency metric. The precuneus is a region often implicated in visual imagery and the recall of memories (Al-Ramadhani et al., 2021; Cavanna and Trimble, 2006; Knauff et al., 2003; Lundstrom et al., 2005; Zhang et al., 2021; Zheng et al., 2023). Similarly, the intraparietal and transverse parietal sulcus are also implicated in visual visual imagery and visual working memory (Bray et al., 2013; Brigadoi et al., 2017; N. Dijkstra et al., 2017; Mellet et al., 2000; Ragni et al., 2020; Slotnick et al., 2005).

For the functional connectome, we found a negative correlation between visual imagery vividness measured with VVIQ scores and global efficiency at the network level.

Global efficiency is indicative of the overall effectiveness of information transfer across the entire network and is measured as the inverse of the average shortest path length from one node to all other nodes in the network (Barbey, 2018; Wang et al., 2018).

The results paint a picture in which a more segregated (higher structural local efficiency and clustering coefficient) and a less globally interconnected network (lower functional global efficiency) at the network level are associated with the ability to experience more vivid visual imagery. Based on our findings, at the network level, it is possible that our results align with what was observed in previous network studies of visual imagery vividness (N. Dijkstra et al., 2017; Milton et al., 2021). These previous findings emphasized the connection between visual areas and frontal regions and a collection of inferior frontal gyrus, visual areas, intraparietal sulcus, and inferior temporal gyrus areas. The fact that these nodes are included in our imagery network suggests that we are in agreement, with findings highlighting how these neural correlates may universally support vivid visual imagery.

This tendency could be because localized networks are better suited to maintain the detailed and specific nature of mental images, as opposed to a globally efficient network that might prioritize overall information transfer over localized detail.

In another study based on the same sample, we observed (in a different frontoparietal control network) that greater musical ability and working memory capacity are correlated with increased global efficiency (Lumaca et al., n.d.). This dovetails with mounting evidence suggesting a link between intelligence and increased global efficiency (Cole et al., 2012; van den Heuvel et al., 2009). It is generally accepted that sustaining cognitive functions relies on a dynamic and context-sensitive balance between functional integration and segregation (Puxeddu et al., 2024; Shine et al., 2018, 2016). Against this backdrop of graph-theoretical dynamics, it is interesting to consider the unique nature of visual imagery vividness as a cognitive phenomenon. Working memory, musicality, and intelligence may require integration across various regions, although such global integration does not seem to support the vividness of visual imagery. We hypothesize that an over-integrated network could lead to a diffusion of the specific neural activity necessary for creating vivid mental images, thereby diluting their clarity and intensity. Moreover, in a highly globally efficient network, neural resources might be distributed more evenly across the network, potentially leading to a less concentrated allocation of resources to the clusters of localized regions that appear crucial for vivid visual imagery.

The findings are also interesting from a developmental perspective as global efficiency changes primarily early in life, but local efficiency continues to change throughout development (Cao et al., 2014; Dennis et al., 2013; Poldrack, 2015; Supekar et al., 2009; Wu et al., 2013) given the reported relationship between age and VVIQ (Floridou et al., 2022; Gulyás et al., 2022).

One avenue for future research could be exploring how the brain’s network structures uniquely support vivid visual imagery as opposed to processing awareness of external visual stimuli and identifying the shared and distinctly neural mechanisms that underlie these forms of conscious experiences (Barnett et al., 2024). Studies indicate a complex interplay in brain networks during conscious experiences. Godwin et al. (2015) showed that awareness of visual stimuli leads to a breakdown in network modularity, suggesting a more integrated brain network during conscious perception of external stimuli (Godwin et al., 2015). In contrast, Sadaghiani et al. (2015) found that a highly modular network structure enhances perceptual efficiency in auditory tasks, highlighting the role of modularity in sensory processing (Sadaghiani et al., 2015). Further, recent research reveals that human visual consciousness involves large-scale networks, including signal detection and arousal/salience networks, attention and executive control networks, with a decrease in the default mode network (Kronemer et al., 2022).

Significant node-level effects in the functional connectome were observed, with the local efficiency in the left lateral occipitotemporal sulcus and the local efficiency and clustering coefficients in the left temporal inferior sulcus showing positive correlations with VVIQ scores, indicating a relationship between these regions’ connectivity and the vividness of visual imagery.

Our hypothesized visual imagery network included the inferior temporal sulcus and gyrus as well as the lateral occipitotemporal sulcus and fusiform gyrus.

The left inferior temporal lobe, particularly the fusiform gyrus, has garnered significant attention in neuroscientific research for its role in visual imagery and vividness (Spagna et al., 2023). This region’s involvement is underscored by a convergence of evidence from various research methodologies, including early work and contemporary work underscoring its importance for domain-general and category-specific visual processing (Haxby et al., 2001; Ishai et al., 2000; Liu et al., 2023) neurological case studies (Bartolomeo, 2002; Thorudottir et al., 2020), effective connectivity analyses (N. Dijkstra et al., 2017), and meta-analytic task fMRI activation studies (Fulford et al., 2018; Spagna et al., 2021).

One previous study that has explored the neural architecture associated with the vividness of visual imagery found increased connectivity between the visual cortex and the frontal cortex in individuals with hyperphantasia suggesting a more integrated network for those with high visual imagery strength (Milton et al., 2021). However, this study categorized participants into high, mid, and low imagery groups without using the full VVIQ scale. Additionally, they used a low-resolution 1.5T MRI scanner and focused primarily on seed-to-voxel connectivity for functional connectivity and fractional anisotropy for structural connectivity (where no differences were found), without exploring the comprehensive structural-functional connectome nor the graph-theoretic properties as performed in our study.

Our research extends the current understanding of visual imagery by demonstrating that local efficiency in two specific regions of the inferior temporal lobe—the left lateral occipitotemporal sulcus and the left temporal inferior sulcus—is positively correlated with the vividness of visual imagery. This finding enriches the existing body of knowledge by highlighting the significance of connectivity in these regions.

While our study did not directly observe effects in the fusiform gyrus per se, the observed local efficiency in the nearby confining regions suggests a potential influence or interaction with the fusiform area (Destrieux et al., 2010).

However, a limitation of our study arises from the coarse resolution of our atlas (Destrieux et al., 2010). This limitation might have restricted our ability to precisely localize the effects within the fusiform gyrus. Additionally, it is crucial to acknowledge that related regions in the left temporal lobe might extend beyond those identified in our fMRI studies. BOLD responses in the anterior temporal lobe are often subject to large magnetic susceptibility artifacts (Wandell and Winawer, 2011), which could potentially obscure the activity in this region.

Another limitation is that our hypothesized network which we defined as apriori is based on task-based fMRI studies. The results we have found do not exclude the possibility that there is another different set of nodes whose structural-functional connectivity organization supports visual imagery vividness. Further work should attempt to find the optimal set of nodes that are maximally predictive or explain the greatest variance in visual imagery vividness potentially using methods that can capture the multivariate nature intrinsic to connectivity measures (Bullmore and Sporns, 2009; Finn et al., 2015; Song et al., 2022).

### Conclusion

Our study demonstrates a compelling link between brain connectivity and visual imagery. Specifically, increased local efficiency and clustering coefficients in the brain’s structural connectome, coupled with decreased global efficiency in the functional connectome, are associated with more vivid visual imagery. Notably, this relationship is further emphasized by significant correlations for local efficiency and clustering coefficient in the left inferior temporal regions, underscoring the intricate interplay between structural and functional brain network dynamics that support the vividness of visual imagery experiences.

## 4. Materials and Methods

### 4.1 Participants

The data were collected under the EU COST Action CA18106. The data is a segment of a larger dataset focused on the Neural Architecture of Consciousness, originating from Aarhus University, Denmark from which other studies have been performed (Fjaeldstad et al., 2022; Lumaca et al., n.d.; Tzioridou et al., 2022). Participants were recruited from the participant pool of the Center of Functionally Integrative Neuroscience at Aarhus University and through local advertisements.

A total of 273 (158 females) participants (out of a total of 300 MRI-scanned participants) were selected for this study based on MRI quality checks. Specifically, their data showed no quality control failures or corruptions in the creation of structural connectomes, nor errors in the functional connectome creation. They were between 18 and 47 years of age (mean=24.79). Detailed sample size considerations for sufficient statistical power are elaborated in Section 1.3.2 of the Technical Annex of the Action, which can be accessed here: https://e-services.cost.eu/files/domain_files/CA/Action_CA18106/mou/CA18106-e.pdf. The study adhered to ethical standards, having received approval from the local ethics committee, De Videnskabsetiske Komitéer for Region Midtjylland in Denmark, and complied with all relevant guidelines and regulations.

### 4.2 Design and Procedures

Participants completed an online questionnaire of the English version of the 32-item VVIQ-2 (Marks, 1995) on a (1-5) Likert scale and were instructed to keep their eyes closed when visualizing the mental image. Online questionnaires were completed approximately one week before completing the MRI scans, which were carried out in one session. One participant scored 1 for all items in the VVIQ and qualified for aphantasia (Blomkvist and Marks, 2023). We ensured that items for this participant were not extreme on other items in the survey.

### 4.3 VVIQ calculation and research tools

We normalized the VVIQ measure to the unit interval, applying a linear transformation so that 0 is the lowest score and 1 is the highest score (Thaler et al., 2014; Zeng et al., 2022). Thus, a summed score X on the scale between 32 and 160 is recorded as VVIQ = (X − 32)/160 in our results.

Plots were performed in the R (4.3.1). We used Shapiro Wilk to test normality as well as tests of uni, bi, and multimodality along with ggplots and raincloud plots to depict correlations and distribution of VVIQ scores using the following R packages (ggplot2, raincloud plots, LaplacesDemon) (Allen et al., 2019; Statisticat, n.d.; Valero-Mora, 2010).

### 4.4 MRI data acquisition

Imaging was performed on a Siemens Magnetom Prisma-fit 3T MRI scanner. The procedure commenced with preliminary scouting scans. This was followed by two sequences of resting-state fMRI, lasting 12 and 6 minutes respectively. The session also included quantitative multi-parameter mapping (approximately 20 minutes) to facilitate the synthetic generation of T1-weighted images. Additionally, high-angular resolution diffusion imaging (HARDI) was conducted over a period of around 10 minutes, all within a single session lasting about one hour. For every participant, we acquired 1500 functional volumes, with a repetition time (TR) of 700 ms and an echo time (TE) of 30 ms. The parameters set were: a voxel size of 2.5 mm^3, a field of view (FOV) of 200 mm, and a flip angle of 53°. The HARDI sequence incorporated multiple diffusion directions: 75 at b = 2500 s/mm^2, 60 at b = 1500 s/mm^2, 21 at b = 1200 s/mm^2, 30 at b = 1000 s/mm^2, 15 at b = 700 s/mm^2, and 10 at b = 5 s/mm^2. These different b-shells were acquired in a single series with a flip angle of 90°, a TR/TE of 2850/71 ms, a voxel size of 2 mm^3, a matrix size of 100 x 100, and 84 slices in total. The primary phase-encoding direction was from anterior to posterior (AP), with an additional acquisition in the opposite phase-encoding direction (PA) at b = xx s/mm^2 for EPI distortion correction.

To create synthetic T1-weighted images, high-resolution longitudinal relaxation rate (R1) and effective proton density (PD) maps were utilized, and obtained through the MPM sequence protocol (Dale et al., 1999; Fischl et al., 1999). Initially, these maps underwent thresholding to align with FreeSurfer’s required units. The R1 map was transformed into a T1 map by inverting its values and applying a zero threshold, followed by a multiplication by 1000. Similarly, the PD map was zero-thresholded and scaled up by a factor of 100. These adjustments were carried out using fslmaths commands. The FreeSurfer’s “mri_synthesize” command was then employed to generate a synthetic FLASH image, using the modified T1 (derived from the adjusted R1 map) and PD maps. Optional arguments were used to enhance the contrast between gray and white matter, with parameters set at 20, 30, and 2.5. In the final step, the synthetic T1-weighted image was reduced by a quarter to meet FreeSurfer’s expected scale (see (Lumaca et al., n.d.) for further details).

### 4.6 DTI processing and structural connectome construction

The preprocessing of the dMRI data was conducted through specialized in-house MATLAB scripts. These scripts effectively filtered out the noise and eliminated common artifacts, including Gibbs ringing, susceptibility distortion, and artifacts caused by eddy currents. Following this, the construction of structural connectomes was carried out utilizing the MRtrix3 software toolkit, with the process entailing a series of intricate steps for each individual participant.

For the connectome generation, each participant’s data underwent a multi-step analysis. Initially, a 5-tissue-type (5tt) image was generated, comprising masks for various brain tissues (cortical and deep grey matter, white matter, CSF, and “other”), crucial for Anatomically-Constrained Tractography (ACT). This was followed by co-registering T1-weighted and DWI images for alignment. Subsequently, a response function for major tissue types (white matter, grey matter, cerebrospinal fluid) was created for each participant. These individual response functions were then combined to form group-level response functions. Fiber Orientation Distributions (FODs) in each brain voxel were calculated using Multi-Shell Multi-Tissue Constrained Spherical Deconvolution (MSMT-CSD). The FODs were then normalized.

We then performed whole-brain probabilistic tractography within the ACT framework, incorporating backtracking. Streamline seeds were dynamically chosen from the FOD image using the SIFT model, with a FOD cutoff of 0.06. The process aimed for a maximum of 1*10^9 streamlines per seed, selecting 10 million streamlines per connectome followed by spherical-deconvolution informed filtering of tractograms (SIFT2). The length of selected streamlines ranged between a maximum of 250mm and a minimum of 20mm. Connectomes were generated using the Destrieux parcellation for cortical areas and FSLs FIRST segmentations for subcortical structures, each scaled by the SIFT proportionality coefficient (mu). The final step involved a custom pipeline for quality control. This process generated JPEGs of the 5TT images and GIFs that allowed fast visual inspection of the data and confirmation of the registration between the T1-weighted image and the B=0 image.

### 4.7 rsfMRI processing and functional connectome construction

The processing of resting-state (rs-fMRI) volumes was implemented by using default surface-based preprocessing routines from the SPM CONN toolbox (version 21a) (Whitfield-Gabrieli (http://www.nitrc.org/projects/conn) (Whitfield-Gabrieli and Nieto-Castanon, 2012), implemented in Matlab (2016b). Functional data was realigned and unwarped without field maps using SPM12 (r7487), employing a 6-parameter transformation for alignment and b-spline interpolation for resampling. Outliers were identified using ART113 based on framewise displacement and global BOLD signal deviations, and an average reference BOLD image was created for each subject excluding all outlier volumes. Coregistration of functional and anatomical data was achieved using mutual information. Functional images were then mapped onto the cortical surface, averaging data across layers between the pial and white matter surfaces. Finally, surface-level functional data were smoothed using 40 iterative diffusion steps. Our denoising process involved a standard pipeline, regressing out confounds like white matter and cerebrospinal fluid (CSF) signals, motion artifacts, outlier scans, session effects, and linear trends. This included the use of CompCor for noise component extraction from white matter and CSF. Bandpass frequency filtering was applied to the BOLD time series to retain frequencies between 0.008 Hz and 0.09 Hz. The effective degrees of freedom of the BOLD signal post-denoising were estimated for all subjects.

We estimated region-to-region connectivity matrices across 22 regions of interest (ROIs) by calculating the functional connectivity strength (Figure 1). This was represented by Fisher-transformed bivariate correlation coefficients derived from a weighted general linear model (GLM) and stored in a functional connectivity matrix. The GLM accounted for associations between BOLD signal time-series of ROI pairs, with weighting to mitigate transient magnetization effects at the start of each run. The connectivity matrix only included Destrieux’s cortical nodes (148x148) (Destrieux et al., 2010). From this matrix, we selected 22 nodes for the hypothesized visual imagery network and two control networks: a frontoparietal network of interest and one occipital control network (16x16) (Fig. 1).

### 4.8 Graph theory analyses

For functional and structural connectivity matrices, the hypothesized visual imagery network, and the frontoparietal and occipital control networks, followed the same pipeline. Connectivity matrices (22×22) and (16×16) were thresholded at a fixed network-level cost range (k) (0<k<1), resulting in binarized, undirected adjacency matrices. The analysis incorporated both positive and negative rs-FC values. To avoid reliance on specific and arbitrary threshold values (e.g., k=0.15) (Langer et al., 2013), graph metrics were aggregated across multiple thresholds (k=0.15–0.30, interval 0.01) (Lumaca et al., 2022). Within this range, brain networks show small-world features (GE and LE have, respectively, larger values than lattice and random graphs of equal size and cost values; Supplementary Fig. S1). From the matrices, four node-level graph theory metrics were computed: clustering coefficient, local efficiency, global efficiency, and betweenness centrality using the Brain Connectivity Toolbox (BCT) (Conti et al., 2019). These metrics offer clear interpretability and are prevalent in network studies. A second-level General Linear Model (GLM) included VVIQ as an explanatory variable, controlling for age, and gender as nuisance regressors. Node-level p-values were adjusted for multiple comparisons using a false discovery rate of q<0.05 (two-tailed), for each graph metric. The four measures were computed at the nodal and global levels. Global metrics are calculated by taking the average of nodal-level measures. At a nodal level: Local efficiency quantifies how well sub-graphs exchange information when the target node is removed. It is the average global efficiency computed on the neighborhood of a node (while removing the target node). Higher local efficiency indicates greater localized information transfer. The clustering coefficient measures the degree of local interconnectedness around a node. For a node, it is the ratio of existing connections between its neighbors divided by the maximum possible connections between its neighbors. Higher clustering suggests dense localized processing. Global efficiency measures how efficiently information can be exchanged between one target node and the rest of the network. It is calculated as the inverse of the average shortest path length between one node and all other nodes in the network. Betweenness centrality captures the influence of a node in exchanging information (hub). It is calculated as the ratio of shortest paths passing through a node divided by all shortest paths in the network. Nodes with high betweenness centrality participate in many shortest paths and can control information transfer.

## Acknowledgments

This article is based upon work from COST Action CA18106 (The Neural Architecture of Consciousness), supported by COST (European Cooperation in Science and Technology). Timo L. Kvamme is supported by an internationalization fellowship from the Carlsberg Foundation. Kristian Sandberg was supported by the Foundation for Research in Neurology and Aarhus University Research Foundation. We thank Katarzyna Hat for her assistance in data collection. We also thank Irene Klærke Mikkelsen, Paola Galdi, and Claude Bajada for providing MRI analysis scripts/processed data available.

## Data availability

Data cannot be shared publicly because it is part of an ongoing study and is thus considered unanonymised under Danish law even if pseudonymized. Researchers who wish to access the data may contact Dr. Kristian Sandberg at The Center of Functionally Integrative Neuroscience and/or The Technology Transfer Office at Aarhus University, Denmark, to make a data sharing contract. After permission has been given by the relevant data committee, data will be made available to the researchers.

## References

Allen, M., Poggiali, D., Whitaker, K., Marshall, T.R., van Langen, J., Kievit, R.A., 2019. Raincloud plots: a multi-platform tool for robust data visualization. Wellcome Open Res 4, 63.

Al-Ramadhani, R.R., Shivamurthy, V.K.N., Elkins, K., Gedela, S., Pedersen, N.P., Kheder, A., 2021. The precuneal cortex: anatomy and seizure semiology. Epileptic Disord. 23, 218–227.

Barbey, A.K., 2018. Network Neuroscience Theory of Human Intelligence. Trends Cogn. Sci. 22, 8–20.

Barnett, B., Andersen, L.M., Fleming, S.M., Dijkstra, N., 2024. Identifying content-invariant neural signatures of perceptual vividness. PNAS Nexus 3, gae061.

Bartolomeo, P., 2002. The relationship between visual perception and visual mental imagery: a reappraisal of the neuropsychological evidence. Cortex 38, 357–378.

Blomkvist, A., Marks, D.F., 2023. Defining and “diagnosing” aphantasia: Condition or individual difference? Cortex. 10.1016/j.cortex.2023.09.004

Bray, S., Arnold, A.E.G.F., Iaria, G., MacQueen, G., 2013. Structural connectivity of visuotopic intraparietal sulcus. Neuroimage 82, 137–145.

Brigadoi, S., Cutini, S., Meconi, F., Castellaro, M., Sessa, P., Marangon, M., Bertoldo, A., Jolicœur, P., Dell’Acqua, R., 2017. On the Role of the Inferior Intraparietal Sulcus in Visual Working Memory for Lateralized Single-feature Objects. J. Cogn. Neurosci. 29, 337–351.

Bullmore, E., Sporns, O., 2009. Complex brain networks: graph theoretical analysis of structural and functional systems. Nat. Rev. Neurosci. 10, 186–198.

Cao, M., Wang, J.-H., Dai, Z.-J., Cao, X.-Y., Jiang, L.-L., Fan, F.-M., Song, X.-W., Xia, M.-R., Shu, N., Dong, Q., Milham, M.P., Castellanos, F.X., Zuo, X.-N., He, Y., 2014. Topological organization of the human brain functional connectome across the lifespan. Dev. Cogn. Neurosci. 7, 76–93.

Cavanna, A.E., Trimble, M.R., 2006. The precuneus: a review of its functional anatomy and behavioural correlates. Brain 129, 564–583.

Cole, M.W., Yarkoni, T., Repovs, G., Anticevic, A., Braver, T.S., 2012. Global connectivity of prefrontal cortex predicts cognitive control and intelligence. J. Neurosci. 32, 8988–8999.

Conti, A., Duggento, A., Guerrisi, M., Passamonti, L., Indovina, I., Toschi, N., 2019. Variability and Reproducibility of Directed and Undirected Functional MRI Connectomes in the Human Brain. Entropy 21. 10.3390/e21070661

Dale, A.M., Fischl, B., Sereno, M.I., 1999. Cortical surface-based analysis. I. Segmentation and surface reconstruction. Neuroimage 9, 179–194.

Dennis, E.L., Jahanshad, N., McMahon, K.L., de Zubicaray, G.I., Martin, N.G., Hickie, I.B., Toga, A.W., Wright, M.J., Thompson, P.M., 2013. Development of brain structural connectivity between ages 12 and 30: a 4-Tesla diffusion imaging study in 439 adolescents and adults. Neuroimage 64, 671–684.

de Pasquale, F., Della Penna, S., Sporns, O., Romani, G.L., Corbetta, M., 2016. A Dynamic Core Network and Global Efficiency in the Resting Human Brain. Cereb. Cortex 26, 4015–4033.

Destrieux, C., Fischl, B., Dale, A., Halgren, E., 2010. Automatic parcellation of human cortical gyri and sulci using standard anatomical nomenclature. Neuroimage 53, 1–15.

Dijkstra, N., 2024. Uncovering the Role of the Early Visual Cortex in Visual Mental Imagery. 10.20944/preprints202402.1684.v1

Dijkstra, N., Bosch, S.E., van Gerven, M.A.J., 2019. Shared Neural Mechanisms of Visual Perception and Imagery. Trends Cogn. Sci. 23, 423–434.

Dijkstra, N., Bosch, S.E., van Gerven, M.A.J., 2017. Vividness of Visual Imagery Depends on the Neural Overlap with Perception in Visual Areas. The Journal of Neuroscience 37, 1367.

Dijkstra, N., Fleming, S.M., 2023. Subjective signal strength distinguishes reality from imagination. Nat. Commun. 14, 1627.

Dijkstra, N., Kok, P., Fleming, S.M., 2022. Perceptual reality monitoring: Neural mechanisms dissociating imagination from reality. Neurosci. Biobehav. Rev. 135, 104557.

Dijkstra, N., Mostert, P., Lange, F.P. de, Bosch, S., van Gerven, M.A., 2018. Differential temporal dynamics during visual imagery and perception. Elife 7. 10.7554/eLife.33904

Dijkstra, N., Zeidman, P., Ondobaka, S., van Gerven, M.A.J., Friston, K., 2017. Distinct top-down and bottom-up brain connectivity during visual perception and imagery. Sci. Rep. 7, 5677.

Finn, E.S., Shen, X., Scheinost, D., Rosenberg, M.D., 2015. Functional connectome fingerprinting: identifying individuals using patterns of brain connectivity. Nature.

Fischl, B., Sereno, M.I., Dale, A.M., 1999. Cortical surface-based analysis. II: Inflation, flattening, and a surface-based coordinate system. Neuroimage 9, 195–207.

Fjaeldstad, A.W., Konieczny, D.T., Fernandes, H., Gaini, L.M., Vejlø, M., Sandberg, K., 2022. The relationship between individual significance of olfaction and measured olfactory function. Current Research in Behavioral Sciences 3, 100076.

Floridou, G.A., Peerdeman, K.J., Schaefer, R.S., 2022. Individual differences in mental imagery in different modalities and levels of intentionality. Mem. Cognit. 50, 29–44.

Fulford, J., Milton, F., Salas, D., Smith, A., Simler, A., Winlove, C., Zeman, A., 2018. The neural correlates of visual imagery vividness - An fMRI study and literature review. Cortex 105, 26–40.

Godwin, D., Barry, R.L., Marois, R., 2015. Breakdown of the brain’s functional network modularity with awareness. Proc. Natl. Acad. Sci. U. S. A. 112, 3799–3804.

Gulyás, E., Gombos, F., Sütöri, S., Lovas, A., Ziman, G., Kovács, I., 2022. Visual imagery vividness declines across the lifespan. Cortex 154, 365–374.

Haxby, J.V., Gobbini, M.I., Furey, M.L., Ishai, A., Schouten, J.L., Pietrini, P., 2001. Distributed and overlapping representations of faces and objects in ventral temporal cortex. Science 293, 2425–2430.

Ishai, A., Ungerleider, L.G., Haxby, J.V., 2000. Distributed neural systems for the generation of visual images. Neuron 28, 979–990.

Iturria-Medina, Y., Sotero, R.C., Canales-Rodríguez, E.J., Alemán-Gómez, Y., Melie-García, L., 2008. Studying the human brain anatomical network via diffusion-weighted MRI and Graph Theory. Neuroimage 40, 1064–1076.

Knauff, M., Fangmeier, T., Ruff, C.C., Johnson-Laird, P.N., 2003. Reasoning, models, and images: behavioral measures and cortical activity. J. Cogn. Neurosci. 15, 559–573.

Kosslyn, S.M., Ganis, G., Thompson, W.L., 2001. Neural foundations of imagery. Nat. Rev. Neurosci. 2, 635–642.

Kronemer, S.I., Aksen, M., Ding, J.Z., Ryu, J.H., Xin, Q., Ding, Z., Prince, J.S., Kwon, H., Khalaf, A., Forman, S., Jin, D.S., Wang, K., Chen, K., Hu, C., Agarwal, A., Saberski, E., Wafa, S.M.A., Morgan, O.P., Wu, J., Christison-Lagay, K.L., Hasulak, N., Morrell, M., Urban, A., Todd Constable, R., Pitts, M., Mark Richardson, R., Crowley, M.J., Blumenfeld, H., 2022. Human visual consciousness involves large scale cortical and subcortical networks independent of task report and eye movement activity. Nat. Commun. 13, 7342.

Langer, N., Pedroni, A., Jäncke, L., 2013. The problem of thresholding in small-world network analysis. PLoS One 8, e53199.

Liu, J., Zhan, M., Hajhajate, D., Spagna, A., Dehaene, S., 2023. Ultra-high field fMRI of visual mental imagery in typical imagers and aphantasic individuals. bioRxiv.

Long, Y., Ouyang, X., Yan, C., Wu, Z., Huang, X., Pu, W., Cao, H., Liu, Z., Palaniyappan, L., 2023. Evaluating test-retest reliability and sex-/age-related effects on temporal clustering coefficient of dynamic functional brain networks. Hum. Brain Mapp. 44, 2191–2208.

Lumaca, M., Keller, P., Baggio, G., Pando, V., Bajada, C.J., Martinez, M.A., Hansen, J.H., Ravignani, R., Joe, N., Vuust, P., Sandberg, K., n.d. Frontoparietal network topology as a neuromarker of music perceptual abilities.

Lumaca, M., Vuust, P., Baggio, G., 2022. Network Analysis of Human Brain Connectivity Reveals Neural Fingerprints of a Compositionality Bias in Signaling Systems. Cereb. Cortex 32, 1704–1720.

Lundstrom, B.N., Ingvar, M., Petersson, K.M., 2005. The role of precuneus and left inferior frontal cortex during source memory episodic retrieval. Neuroimage 27, 824–834.

Marks, D.F., 1995. New directions for mental imagery research. Journal of Mental Imagery 19, 153–167.

Mellet, E., Tzourio-Mazoyer, N., Bricogne, S., Mazoyer, B., Kosslyn, S.M., Denis, M., 2000. Functional anatomy of high-resolution visual mental imagery. J. Cogn. Neurosci. 12, 98–109.

Milton, F., Fulford, J., Dance, C., Gaddum, J., Heuerman-Williamson, B., Jones, K., Knight, K.F., MacKisack, M., Winlove, C., Zeman, A., 2021. Behavioral and Neural Signatures of Visual Imagery Vividness Extremes: Aphantasia versus Hyperphantasia. Cereb Cortex Commun 2, tgab035.

Monti, M.M., Vanhaudenhuyse, A., Coleman, M.R., Boly, M., Pickard, J.D., Tshibanda, L., Owen, A.M., Laureys, S., 2010. Willful modulation of brain activity in disorders of consciousness. N. Engl. J. Med. 362, 579–589.

Owen, A.M., Coleman, M.R., Boly, M., Davis, M.H., Laureys, S., Pickard, J.D., 2006. Detecting awareness in the vegetative state. Science 313, 1402.

Pearson, J., 2019. The human imagination: the cognitive neuroscience of visual mental imagery. Nat. Rev. Neurosci. 20, 624–634.

Pearson, J., Naselaris, T., Holmes, E.A., Kosslyn, S.M., 2015. Mental Imagery: Functional Mechanisms and Clinical Applications. Trends Cogn. Sci. 19, 590–602.

Poldrack, R.A., 2015. Is “efficiency” a useful concept in cognitive neuroscience? Dev. Cogn. Neurosci. 11, 12–17.

Puxeddu, M.G., Faskowitz, J., Seguin, C., Yovel, Y., Assaf, Y., Betzel, R., Sporns, O., 2024. Relation of connectome topology to brain volume across 103 mammalian species. PLoS Biol. 22, e3002489.

Ragni, F., Tucciarelli, R., Andersson, P., Lingnau, A., 2020. Decoding stimulus identity in occipital, parietal and inferotemporal cortices during visual mental imagery. Cortex 127, 371–387.

Sadaghiani, S., Poline, J.-B., Kleinschmidt, A., D’Esposito, M., 2015. Ongoing dynamics in large-scale functional connectivity predict perception. Proc. Natl. Acad. Sci. U. S. A. 112, 8463–8468.

Shine, J.M., Aburn, M.J., Breakspear, M., Poldrack, R.A., 2018. The modulation of neural gain facilitates a transition between functional segregation and integration in the brain. Elife 7. 10.7554/eLife.31130

Shine, J.M., Bissett, P.G., Bell, P.T., Koyejo, O., Balsters, J.H., Gorgolewski, K.J., Moodie, C.A., Poldrack, R.A., 2016. The Dynamics of Functional Brain Networks: Integrated Network States during Cognitive Task Performance. Neuron 92, 544–554.

Skottnik, L., Linden, D.E.J., 2019. Mental Imagery and Brain Regulation-New Links Between Psychotherapy and Neuroscience. Front. Psychiatry 10, 779.

Slotnick, S.D., Thompson, W.L., Kosslyn, S.M., 2005. Visual mental imagery induces retinotopically organized activation of early visual areas. Cereb. Cortex 15, 1570–1583.

Song, C., Sandberg, K., Rutiku, R., Kanai, R., 2022. Linking human behaviour to brain structure: further challenges and possible solutions. Nat. Rev. Neurosci. 10.1038/s41583-022-00614-4

Spagna, A., Hajhajate, D., Liu, J., Bartolomeo, P., 2021. Visual mental imagery engages the left fusiform gyrus, but not the early visual cortex: A meta-analysis of neuroimaging evidence. Neurosci. Biobehav. Rev. 122, 201–217.

Spagna, A., Heidenry, Z., Miselevich, M., Lambert, C., Eisenstadt, B.E., Tremblay, L., Liu, Z., Liu, J., Bartolomeo, P., 2023. Visual mental imagery: evidence for a heterarchical neural architecture. Phys. Life Rev. 10.1016/j.plrev.2023.12.012

Sporns, O., 2011. The human connectome: a complex network. Ann. N. Y. Acad. Sci. 1224, 109–125.

Statisticat, L.L.C., n.d. LaplacesDemon: Complete environment for Bayesian inference. R package version.

Supekar, K., Musen, M., Menon, V., 2009. Development of large-scale functional brain networks in children. PLoS Biol. 7, e1000157.

Thaler, L., Wilson, R.C., Gee, B.K., 2014. Correlation between vividness of visual imagery and echolocation ability in sighted, echo-naïve people. Exp. Brain Res. 232, 1915–1925.

Thorudottir, S., Sigurdardottir, H.M., Rice, G.E., Kerry, S.J., Robotham, R.J., Leff, A.P., Starrfelt, R., 2020. The architect who lost the ability to imagine: The cerebral basis of visual imagery. Brain Sci. 10, 59.

Tullo, M.G., Almgren, H., Van de Steen, F., Sulpizio, V., Marinazzo, D., Galati, G., 2022. Individual differences in mental imagery modulate effective connectivity of scene-selective regions during resting state. Brain Struct. Funct. 227, 1831–1842.

Tzioridou, S., Dresler, M., Sandberg, K., Mueller, E.M., 2022. The role of mindful acceptance and lucid dreaming in nightmare frequency and distress. Sci. Rep. 12, 15737.

Valero-Mora, P.M., 2010. ggplot2: Elegant Graphics for Data Analysis. Springer.

van den Heuvel, M.P., Stam, C.J., Kahn, R.S., Hulshoff Pol, H.E., 2009. Efficiency of functional brain networks and intellectual performance. J. Neurosci. 29, 7619–7624.

Wandell, B.A., Winawer, J., 2011. Imaging retinotopic maps in the human brain. Vision Res.

Wang, B., Li, P., Li, D., Niu, Y., Yan, T., Li, T., Cao, R., Yan, P., Guo, Y., Yang, W., Ren, Y., Li, X., Wang, F., Yan, T., Wu, J., Zhang, H., Xiang, J., 2018. Increased Functional Brain Network Efficiency During Audiovisual Temporal Asynchrony Integration Task in Aging. Front. Aging Neurosci. 10, 316.

Whitfield-Gabrieli, S., Nieto-Castanon, A., 2012. Conn: a functional connectivity toolbox for correlated and anticorrelated brain networks. Brain Connect. 2, 125–141.

Wig, G.S., 2017. Segregated Systems of Human Brain Networks. Trends Cogn. Sci. 21, 981–996.

Winlove, C.I.P., Milton, F., Ranson, J., Fulford, J., MacKisack, M., Macpherson, F., Zeman, A., 2018. The neural correlates of visual imagery: A co-ordinate-based meta-analysis. Cortex 105, 4–25.

Wu, K., Taki, Y., Sato, K., Hashizume, H., Sassa, Y., Takeuchi, H., Thyreau, B., He, Y., Evans, A.C., Li, X., Kawashima, R., Fukuda, H., 2013. Topological organization of functional brain networks in healthy children: differences in relation to age, sex, and intelligence. PLoS One 8, e55347.

Zeng, C., Fielding, D., Peeters, R., Wesselbaum, D., 2022. Visual imagery skills and risk attitude. Sci. Rep. 12, 21415.

Zhang, M., Duan, F., Wang, S., Zhang, K., Chen, X., Sun, Z., 2021. A Study of the Brain Network Connectivity in Visual-Word Pairing Associative Learning and Episodic Memory Reactivating Task. Comput. Intell. Neurosci. 2021, 5579888.

Zheng, Y., Xie, L., Huang, Z., Peng, J., Huang, S., Guo, R., Huang, J., Lin, Z., Zhuang, Z., Yin, J., Hou, Z., Ma, S., 2023. Enhanced activity of the left precuneus as a predictor of visuospatial dysfunction correlates with disease activity in rheumatoid arthritis. Eur. J. Med. Res. 28, 276.

