## Supplementary figures and images for "Vividness of Visual Imagery Supported by Intrinsic Structural-Functional Brain Network Dynamics"

### supplementary figure s1

**Supplementary Figure S1**

Structural connectome

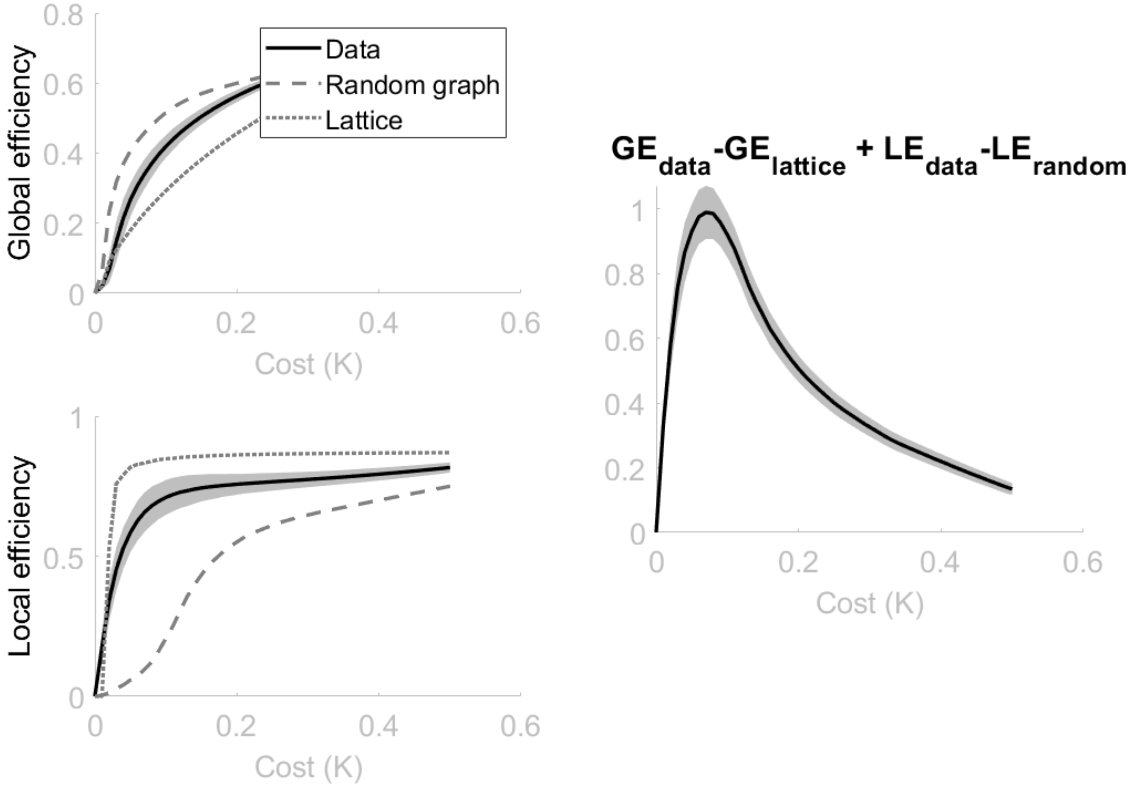


Functional connectome

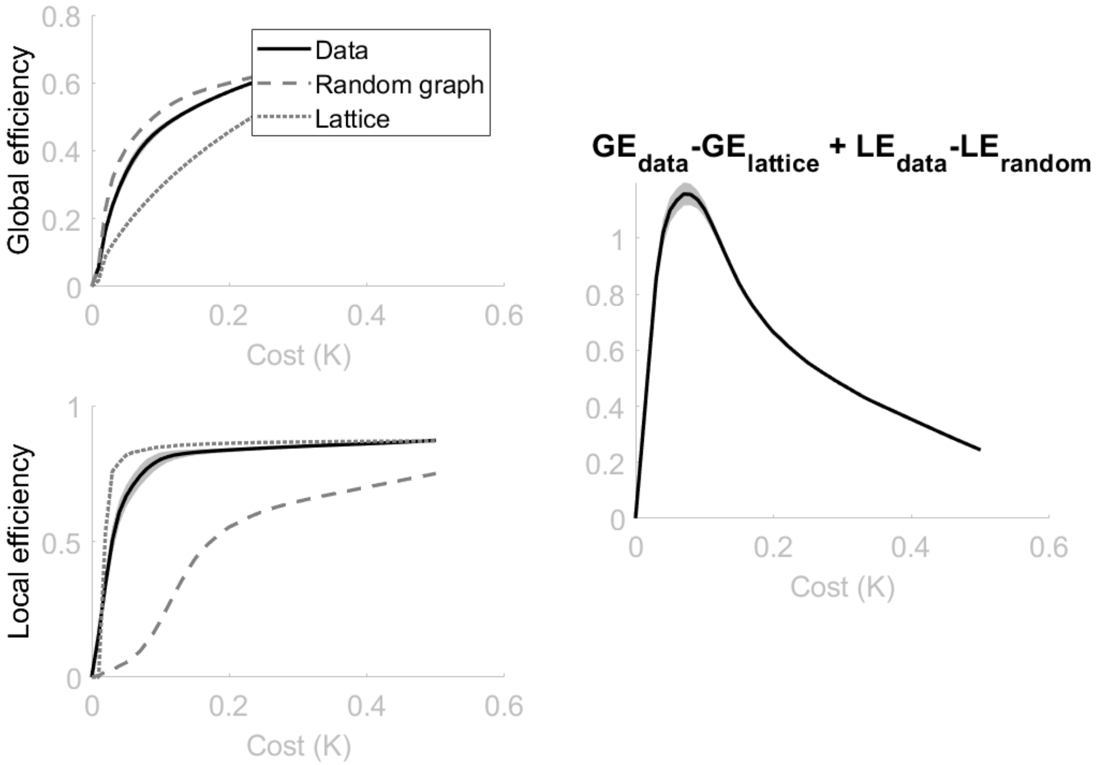
